# Engineered inter-species amino acid cross-feeding increases population evenness in a synthetic bacterial consortium

**DOI:** 10.1101/426171

**Authors:** Marika Ziesack, Travis Gibson, John K.W. Oliver, Andrew M. Shumaker, Bryan B. Hsu, David T. Riglar, Tobias W. Giessen, Nicholas V. DiBenedetto, Lynn Bry, Jeffrey C. Way, Pamela A. Silver, Georg K. Gerber

## Abstract

In nature, microbes interact antagonistically, neutrally or beneficially. To shed light on the effects of positive interactions in microbial consortia we introduced metabolic dependencies and metabolite overproduction into four bacterial species. While antagonistic interactions govern the wildtype consortium behavior, the genetic modifications alleviated antagonistic interactions and resulted in beneficial interactions. Engineered cross-feeding increased population evenness, a component of ecological diversity, in different environments including in a more complex gnotobiotic mouse gut environment. Our findings suggest that metabolite cross-feeding could be used as a tool for intentionally shaping microbial consortia in complex environments.

**Importance:** Microbial communities are ubiquitous in nature. Bacterial consortia live in and on our body and in our environment and more recently, biotechnology is applying microbial consortia for bioproduction. As part of our body, bacterial consortia influence us in health and disease. Microbial consortia function is determined by its composition, which in turn is driven by the interactions between species. Further understanding of microbial interactions will help us deciphering how consortia function in complex environments and may enable us to modify microbial consortia for health and environmental benefits.

## Introduction

In nature, microbes occur as conglomerates of various species with diverse sets of genomes and metabolic capabilities (1). Composition of these consortia arises from networks of ecological interactions and determines function and performance in varying environments. Ecological interactions can be antagonistic (e.g. competition for nutrients), neutral (no interaction) or beneficial (e.g. resource sharing) (2). Studies suggest that both antagonistic and beneficial interactions drive microbial consortia composition (3). According to classic evolutionary theory, competitive interactions favoring survival of one species over another should be prevalent, and a body of experimental research supports that hypothesis (4–6). Recent studies of the microbiota also highlight the importance of beneficial interactions, particularly those that provide “public goods” (7, 8). “Public goods” are metabolites that are released into the environment, and not only benefit the producer but also neighboring cells.

A meta-analysis of 77 studies revealed that cross-feeding of metabolites between naturally occurring bacteria is common (7). Metabolite cross-feeding may be explained by comparative advantages of metabolic capabilities in some species over others, and removal of biochemical conflicts in biosynthetic pathways within cells (9, 10). The evolution of such mutually beneficial interactions may occur as follows. Metabolic leakiness in strain A can serve as the first step towards metabolic dependencies as it results in products leaking into the environment where neighboring strain B can take them up (11, 12). Strain B may lose the ability to make the metabolites itself, resulting in auxotrophy. Indeed, microbial auxotrophies are common in natural communities. In a study of 949 eubacterial genomes, 76% were found to lack at least one of 25 different metabolic biosynthesis pathways (13).

There is increasing recognition that maintaining species diversity – a measure of richness and evenness within a population – in microbiomes is important to ecosystem health (14). A study that analyzed composition of a syntrophic consortium of methanotrophs found that amino acid cross-feeding may help with retaining species diversity which leads to higher robustness with regard to limiting loss or dominance of species in the consortium (15). Despite the fact that bacteria competed for similar nutritional niches, the consortium composition remained stable. Thus, metabolite crossfeeding could serve as a mechanism to preserve species diversity by limiting the ability of some consortium members to dominate and eliminate other species.

Amino acid cross-feeding is also an attractive means to introduce cooperation into synthetic microbial consortia. Numerous studies have engineered pairwise amino acid cross-feeding in *E. coli* with native amino acid production levels and generated quantitative models to describe their behavior (16–21). Most prior synthetic biology studies that have engineered cooperativity in bacterial communities have used a single species (21–24). However, natural microbial ecosystems contain a diversity of interacting species.

In advancing synthetic biology to real applications in complex environments, it will be useful to expand engineering capabilities to diverse, multi-species consortia. Importantly, bacteria from naturally occurring ecosystems are likely to have pre-existing interactions, which are often competitive. In a given bacterial consortium that is dominated by antagonistic interactions, synthetically introduced positive interactions need to overcome these antagonistic interactions in order to cause a net beneficial effect. Here we engineer metabolite cross-feeding in a synthetic consortium of four different bacterial species and demonstrate net beneficial behavior. We further observe increased population evenness–a component of species diversity within a population– in different environments *in vitro* and *in vivo.*

## Results

### Engineering of metabolic dependency and overproduction between four species

For our consortium, we chose four bacterial species that naturally reside in the mammalian gut, a complex environment of intense interest both from the basic scientific and biomedical perspectives (Figure 1a). The organisms were chosen in part for their propensity to colonize in the colon, which we hypothesized could allow for sufficient cross-feeding opportunities. A second crucial criteria for the selected organisms was the availability of tools for efficient genetic manipulation; to date, a very small subset of commensal gut microbes are genetically tenable and thus a wider range of organisms was not feasible. Nonetheless, these organisms represent gut residents with different spatial niches and carbohydrate utilization preferences: *E. coli* and S. Typhimurium tend to utilize simpler carbohydrates as found in the gut lumen (25, 26), whereas *Bacteroides spp.* tend to be mucosally associated and utilize more complex carbohydrates (27).

**Figure 1.**
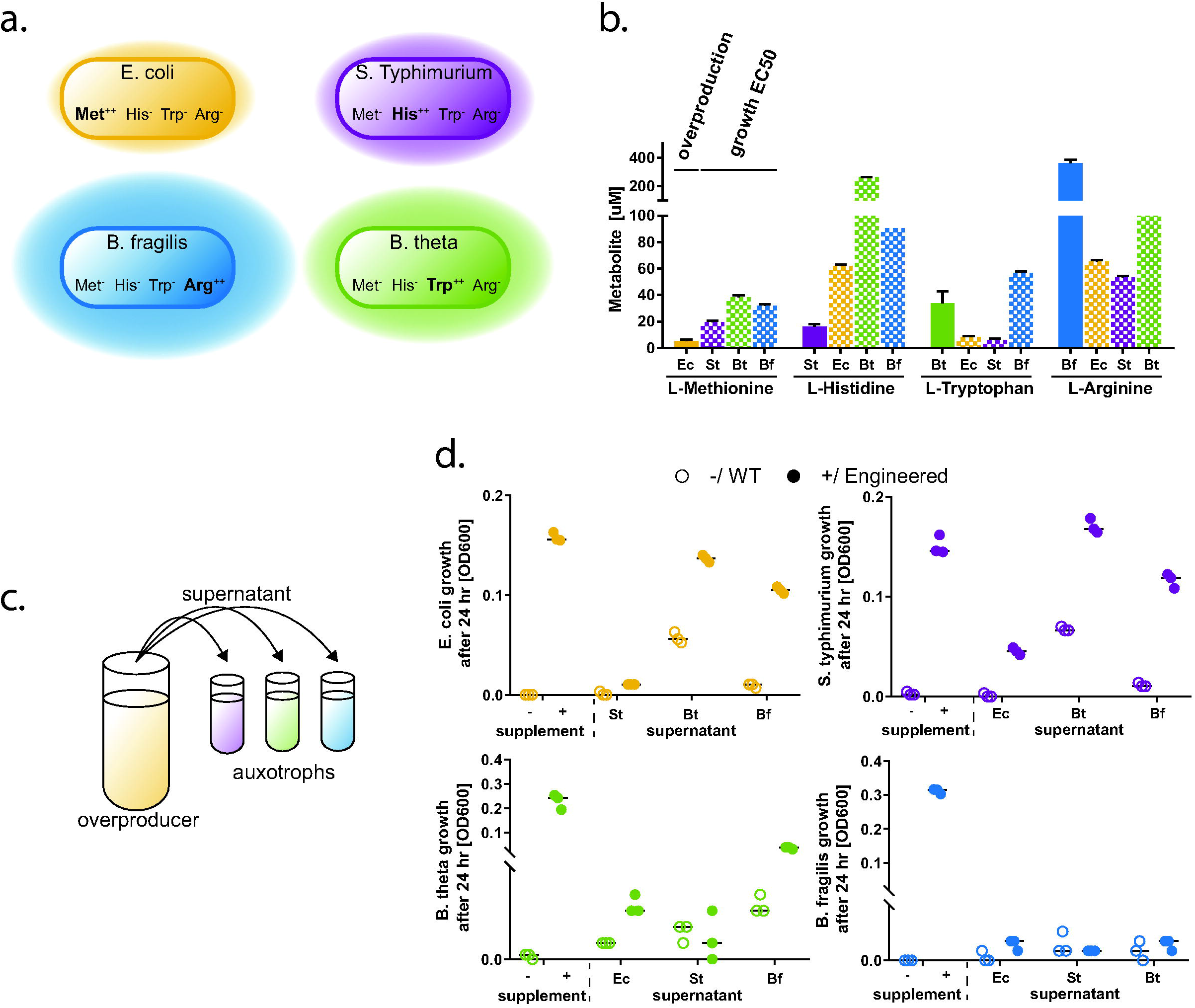
Engineered Metabolite Overproduction and Growth Requirements. **a.** Consortia design. Each strain is auxotrophic for three amino acids and overproduces one. **b.** Quantification of overproduction and growth requirement (EC50). Overproduction in each engineered strain was measured via LC-MS with appropriate amino acid standard. Metabolite growth requirement EC50 was measured in media with full supplementation of two amino acids and varying amounts of one other. **c.** Cross-feeding capabilities of each strain were assessed by testing for rescue of auxotroph growth in supernatants obtained from overproducers. **d.** Growth assay of auxotrophic strains. Auxotroph strains were grown in media without and with addition of all cross-fed amino acids (left side of graphs, −/+) which served as a reference point. Each strain was also grown in fresh media without supplementation and supernatant of engineered overproducers or WT counterpart at a 1:1 ratio. Shown are three biological replicates with median indicated as horizontal lines.

*Escherichia coli* NGF-1 was originally isolated from BALB/c mice, has been shown to stably colonize the mouse gut, and can be engineered with standard genetic tools (28–30). *Salmonella enterica* serovar Typhimurium LT2 *(S.* Typhimurium), an attenuated gut pathogen, was rendered further harmless by removing the pathogenicity islands SPI1 and SPI2, and thus causes no disease when administered to mice (31). The two *Bacteroides* species, *Bacteroides thetaiotaomicron* and *Bacteroides fragilis,* are human commensals that can achieve high abundance in the mammalian gut, and are also genetically tractable.

We engineered each of the constituent species to depend on three of the four metabolites L-methionine, L-histidine, L-tryptophan and L-arginine (hereafter referred to as Met, His, Trp and Arg) and to overproduce the remaining amino acid (Figure 1a). As we noted, amino acid cross-feeding has been described as a prominent natural mechanism (7) and has previously been used extensively in single species engineering efforts. Our rationale for choosing these specific amino acids for cross-feeding was in part based on feasibility of engineering overproduction and auxotrophies in the selected species based on prior work. Additionally, we considered the amino acid composition of proteins across a variety of bacterial taxa, which indicated lower use of Met, His and Trp, and higher for Arg (32). We therefore hypothesized that these particular amino acids would offer a range of nutrient requirements for testing and evaluating consortium behavior.

The auxotrophies introduced for each strain were: *E. coli:* His, Trp and Arg; S.Typhimurium: Met, Trp, Arg; *B. thetaiotaomicron:* Met, His, Arg; *B.fragilis:* Met, His, Trp. *E. coli* and S. Typhimurium were engineered by sequential phage transduction from three single auxotroph strains. *E. coli* was transduced with genome fragments from BW25113 that contained insertions in argA, trpC, hisA (see Methods) and S. Typhimurium with genome fragments of the same parent strain with insertions in argA, trpC, metA. *Bacteroides spp.* triple knockout generation utilized the pExchange-tdk vector to precisely delete *metA, hisG* and *argF* in *B. thetaiotaomicron* and *metA, hisG* and *trpC* in *B. fragilis.* We introduced the ability to overproduce one amino acid in each strain by using anti-metabolite selection methods *(E. coli:* Met; S. Typhimurium: His; *B. thetaiotaomicron:* Trp; *B. fragilis:* Arg) and identified putatively causal mutations by sequencing and SNP analysis (Table 1, Methods).

**Table 1.**
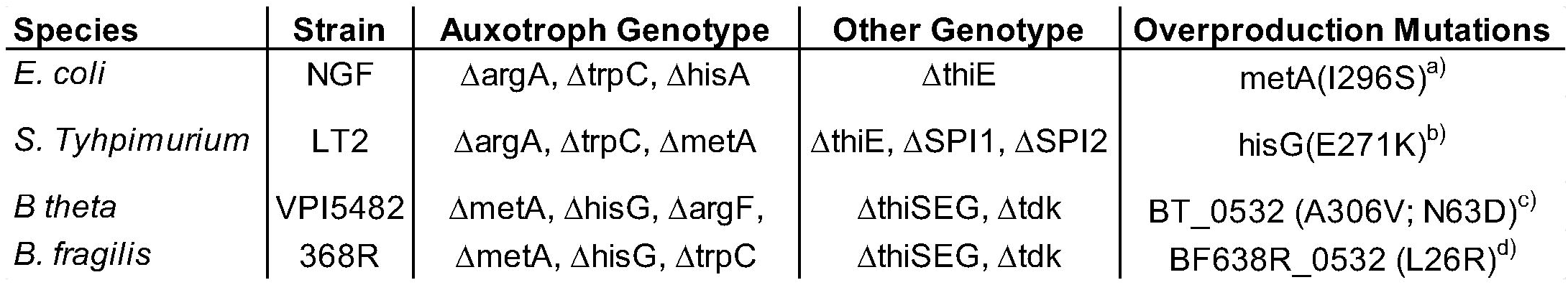
Engineered strains, engineered genotypes and a subset of identified SNPs previously implicated as causal mutations. ^a)^prevents feedback inhibition (45), ^b)^decouples from histidine feedback inhibition (46); ^c)^trpE, removes feedback inhibition (47); ^d)^arginine repressor, nonfunctional (48)

To assess the auxotrophic strains’ requirements for the metabolite, we measured growth on varying concentrations of each metabolite in the presence of non-limiting concentrations of all the other metabolites (Figure S1). We found that each strain has distinct requirements for the amino acids that it was auxotrophic for. In general, the *Bacteroides spp.* have higher amino acid requirements compared to *E. coli* and S. Typhimurium, which could in part be due to differences in their capability to transport/uptake amino acids. We note also that the observed amino acid requirements correlate with differing amino acid compositions in proteins of these species (33).

Overproduction of metabolites was measured in comparison to a defined amino acid standard using LC-MS (Figure 1b, filled bars, Figure S6). In order to compare overproduction with each species’ amino acid requirements, we fit a sigmoidal curve to the growth response data and calculated the EC50 requirement (Figure S1, and 1b, shaped bars). In this context, the EC50 requirement describes the concentration of supplemented metabolite that allows for half-maximum growth of the respective strain. These data on amino acid requirements and overproduction levels provide information about the expected relative strengths of the engineered beneficial interactions.

Engineered *B. fragilis* overproduced Arg at 362 uM - a concentration exceeding EC50 requirements of any of the other strains. Corresponding supplementation allows for more than half-maximum growth of either one of the strains. *B. thetaiotaomicron* overproduced Trp at 34 uM, which exceeded the EC50 requirements of *E. coli* and S. Typhimurium but not *B. fragilis.* Overproduction from *E. coli* (Met at 5.2 uM) and S. Typhimurium (His at 16 uM) were lower than any of the EC50 requirement values. EC50 requirement values exceeded *E. coli* Met overproduction by 4-, 7-, and 6-fold and S. Typhimurium His overproduction by 12-, 51- and 17-fold. Overall, these findings suggest relatively strong cross-feeding from *Bacteroides spp,* moderate cross-feeding from *E. coli,* and the weakest cross-feeding from S. Typhimurium to other strains. Note that WT *E. coli* and S. Typhimurium do not produce detectable levels of the respective amino acids in supernatant, whereas WT *B. theta* produces 5.7 ± 0.8 μM Trp and WT *B. fragilis* produces 1.7 ± 1.2 μM Arg (Figure S6).

To provide further insight into the ability of each overproducer strain to support the auxotrophic strains, we performed assays in which each auxotrophic strain was grown in the presence of supernatant from the overproducer strain or its WT counterpart (Figure 1c). Overall, our overproduction and requirement assay described above predicted growth in the supernatant assay. The majority of WT strain supernatants supported no to minimal growth of the auxotrophs, whereas the overproducer supernatants variably supported growth depending on the match between overproduction and metabolite requirements of the respective pair of strains (Figure 1d). Exceptions to this observation were WT *B. theta* supernatant, which supported some growth of *E. coli* and S. Typhimurium, and WT *B. fragilis* supernatant, which supported some growth of *B. theta.* This is consistent with our finding that the WT strains of these organisms produced a small amount of the respective amino acid (Figure S6). Growth of *E. coli* was rescued by supernatant from overproducing *B. theta* and *B. fragilis* but not S. Typhimurium. S. Typhimurium growth was rescued by all overproducer supernatants. *B. theta* growth was partially rescued by *E. coli* and *B. fragilis* but not S. Typhimurium overproducer supernatant. *B. fragilis* growth was not rescued by any of the strains’ overproducer supernatants but is rescued in co-culture (see below).

### Engineered cross-feeding introduces beneficial interactions in the consortium

To characterize the behavior and interaction structure of the consortium as a whole, we performed a set of experiments in which we cultured all members together while varying the starting inoculum of one member at a time. By measuring the growth trajectories of all strains simultaneously, we can analyze how fluctuations in the growth of one member of the consortium influences changes in the growth of other members over time, and thus infer the interactions that are occurring. We performed these experiments under five conditions for both WT and engineered consortia.

In condition 1, which served as the baseline, we inoculated all consortium members at equal ratios (Figure 2 a, b), and in conditions 2-5 we reduced the starting inoculum of each of the four species in turn by a factor of 10 (Figure 2 c,d). We grew the consortia anaerobically at 37 °C for 27 hours in M9 minimal media with specific modifications as described in Methods, without supplementation of any of the cross-fed amino acids, and with 0.5% starch and 0.5% glucose as carbon sources. We assessed growth of each strain in co-culture over time via strain-specific qPCR.

**Figure 2.**
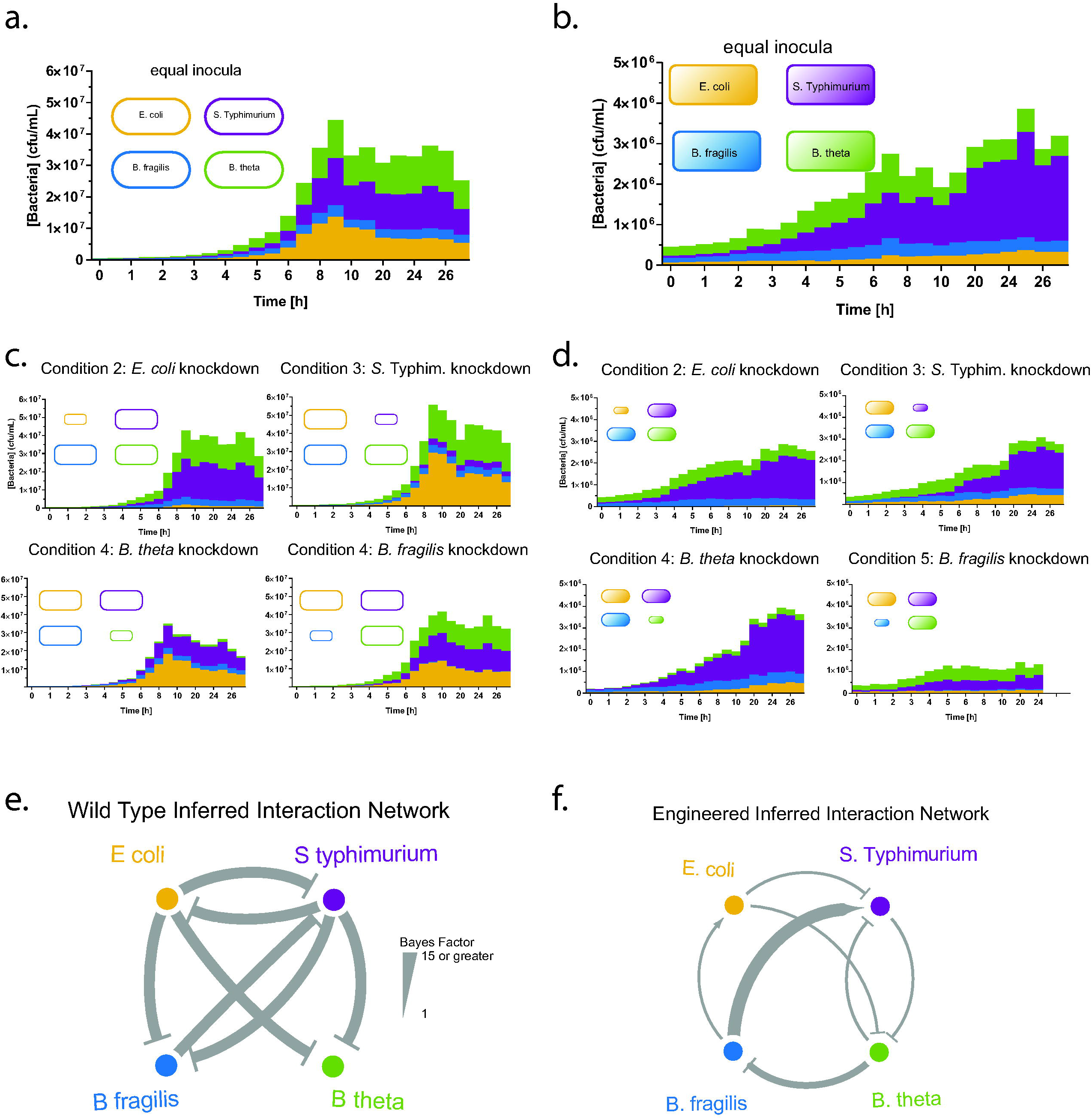
Engineered consortium exhibits added beneficial interactions and reduced antagonistic interactions. **a./b.** Growth trajectories in anaerobic batch co-cultures for WT (**a**.) and engineered (b.) consortia. All strains were inoculated at equal ratios and grown anaerobically at 37 °C for 27 hours in M9 minimal media with specific modifications as described in Methods, without supplementation of any of the cross-fed amino acids, and with 0.5% starch and 0.5% glucose as carbon sources. The engineered consortium grows to an order of magnitude lower density than the WT counterpart. **c./d**. Growth trajectories of consortia with each strain inoculum reduced by 10-fold of WT (**c**.) and engineered (**d**.) consortia. These data combined were used to infer a network interaction using the MDSINE algorithm. **e./f**. Inferred interaction networks for WT (**e**.) and engineered (**f**.) consortia. WT consortia show mainly negative interactions with large Bayes factors. Engineered consortia’s negative interactions have reduced Bayes factors and positive interactions with strong Bayes factors occurred between *B. fragilis* and *E. coli* and S. Typhimurium.

We were able to record growth of each strain in WT consortia indicating that conditions were permissive for each strain (Figure 2a). We also recorded growth of each strain in the engineered consortia, which indicates that cross-feeding capabilities were sufficient to support growth (Figure 2b). As expected based on our previous experiment the engineered consortia did not fully rescue all of the auxotrophies and hence grew to about 10-fold lower overall cfu/mL. In the WT consortium, knockdown of one species persists throughout culturing, whereas in the engineered consortium S. Tyhpimurium and *B. theta* are able to recover to baseline ratios (Figure 2 c,d). Two further characteristics of the engineered consortia are notable. In all five conditions, S. Typhimurium appears to dominate the culture, and *B. fragilis* knockdown causes a decrease in overall consortia growth by 2-fold.

We note that no *B. fragilis* growth in supernatant was observed, whereas we do see growth in the co-culture setting. This difference may be due to several reasons. One possibility is that in the co-culture setting *B. fragilis* may have time to grow to sufficient biomass such that the growth inhibition is blunted, whereas in the supernatant setting the other organisms can accumulate high concentrations of substances that inhibit initial *B. fragilis* growth. Alternately, other consortium members may be able to actively buffer negative effects. For example, we have observed that S. Typhimurium acidifies the media, which, without appropriate buffering, strongly decreased *Bacteroides spp.* growth (data not shown).

To visualize how lowering the inoculum of each strain (“knockdown”) affected growth of the other strains, we plotted log-fold ratios (logR) comparing the “knockdown” condition to the equal inoculum condition (Figure S2, S3). Positive logR values for strain A indicate more growth of that strain in the absence of the “knockdown” strain B, suggesting a net antagonistic effect of strain B on strain A; conversely, a negative logR value suggests a net positive effect. We see either negligible or positive logR values for the WT consortium (Figure S2), suggesting that it is dominated by antagonistic interactions. In contrast, for the engineered consortium we see negligible or negative logR values (Figure S3), suggesting antagonistic interactions have been reduced and positive interactions were introduced.

To quantitate interactions present in the consortium, taking into account both direct and cascading effects, we used a statistical method based on well-established techniques for inferring microbial interaction networks from time-series data (33–36). Briefly, the method models the rate of change in growth of species A as a function of the concentrations of itself and all other species present in the ecosystem. (See the Methods section for further details.) Applying our inference method to the data presented in Figure 2 a-d, we obtained a network of inferred interactions and accompanying estimates of confidence in each edge in the network (Figure 2 e, f, Supplemental Figure S4). Consistent with our qualitative visualization, all inferred interactions for the WT consortium were antagonistic (Figure 2 e). In contrast, we see two inferred positive interactions in the engineered consortium network, from *B. fragilis* to *E. coli* and S. Typhimurium with high inference confidence, which is also consistent with our qualitative visualization (Figure 2 f). Further, although some inferred negative interactions are still present in the engineered consortium network, the confidence in these predictions is much lower than in the WT network, indicating that the model finds greatly reduced evidence for antagonistic interactions in the engineered consortium. Taken together, our findings indicate that introducing metabolic dependencies and overproduction capacities alleviated antagonistic interactions and introduced some beneficial interactions in the consortia.

### Engineered consortia show higher population evenness than WT counterpart *in vitro*

To gain insight into how beneficial interactions affect consortia behavior, we compared consortia compositions in environments with different nutrient levels (Figure 3). We grew consortia in batch culture for 24 hr anaerobically in modified M9 media as described in Methods with and without supplementation of 1 mM of the cross-fed amino acids and 0.2% starch and 0.5% glucose as carbon sources. As described in the previous trajectory experiment the engineered consortia were able to grow without amino acid supplementation albeit to lower capacity compared to its WT counterpart. Addition of cross-fed amino acids rescued growth of the engineered strains to levels equivalent of the WT counterparts. This is in accordance with previously described consortia characterization (Figure 1) which shows that not all overproduction levels were sufficient to fully rescue growth of all species.

**Figure 3.**
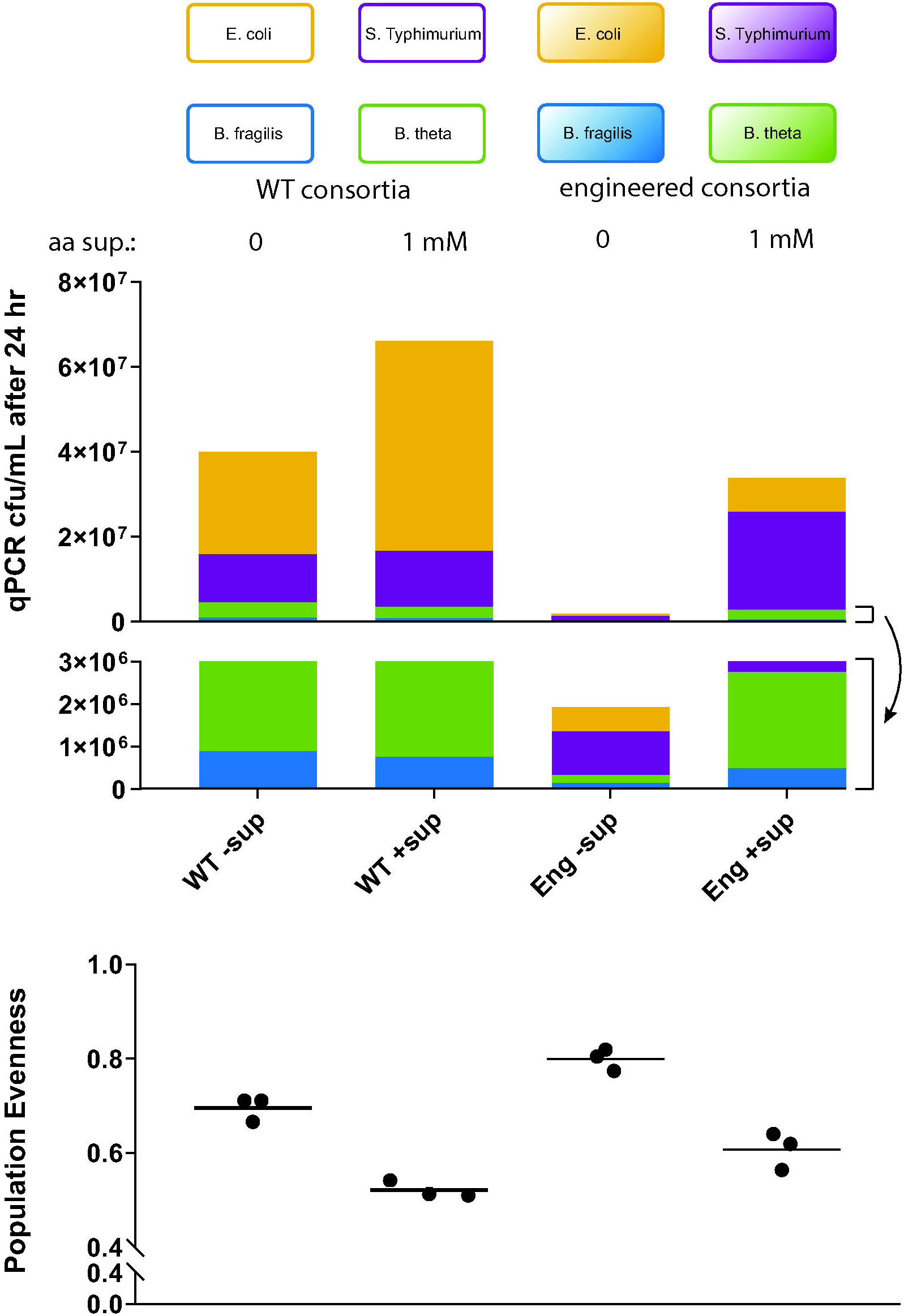
Engineered Consortia exhibit increased population evenness *in vitro.* WT and engineered consortia were grown anaerobically in batch co-culture for 24 hr anaerobically in modified M9 media as described in Methods with and without supplementation of 1 mM of the cross-fed amino acids and 0.2% starch and 0.5% glucose as carbon sources. Upper panel: average population composition after 24 hr. Lower panel: Quantified Pielou Evenness. The engineered consortium has higher population evenness with or without amino acid supplementation compared to WT consortium.

An important ecological measure of consortium behavior is its population evenness in different environments, which is frequently computed using Pielou evenness index. This measure indicates the extent to which abundances of different species in an ecosystem are even or balanced; a value of 1.0 indicates equal abundances of all species, whereas lower values indicate an ecosystem dominated by a subset of the total species present, and a value of 0 indicates one species completely dominates the community.

The highest population evenness occurred in the engineered consortia without amino acid supplementation, a setup in which cross-feeding via engineered overproduction and auxotrophies was expected to be maximized. The WT consortium also exhibited a similar trend, showing higher population evenness when amino acid supplementation was omitted. However, the population evenness attained by the WT consortium in this condition was still considerably less than that of the engineered consortium in the same condition. Overall, our results suggest that introduction of cross-feeding can increase population evenness in conditions favoring these beneficial interactions.

### Engineered consortia show higher population evenness than WT counterpart in the mouse gut

We investigated the behavior of our consortium in the mammalian gut, using gnotobiotic mice as a controlled yet relatively complex environment for evaluation. To investigate the role of amino acid cross-feeding *in vivo,* we altered amino acid levels in the gut by changing the animal’s diet (37). Four groups of five germfree mice were fed standard or low protein (3%) chow, and gavaged with either the WT or engineered consortium (Figure 4). The consortia were allowed to colonize for 10 days, and then stool samples were collected and interrogated via qPCR with species-specific primers.

**Figure 4.**
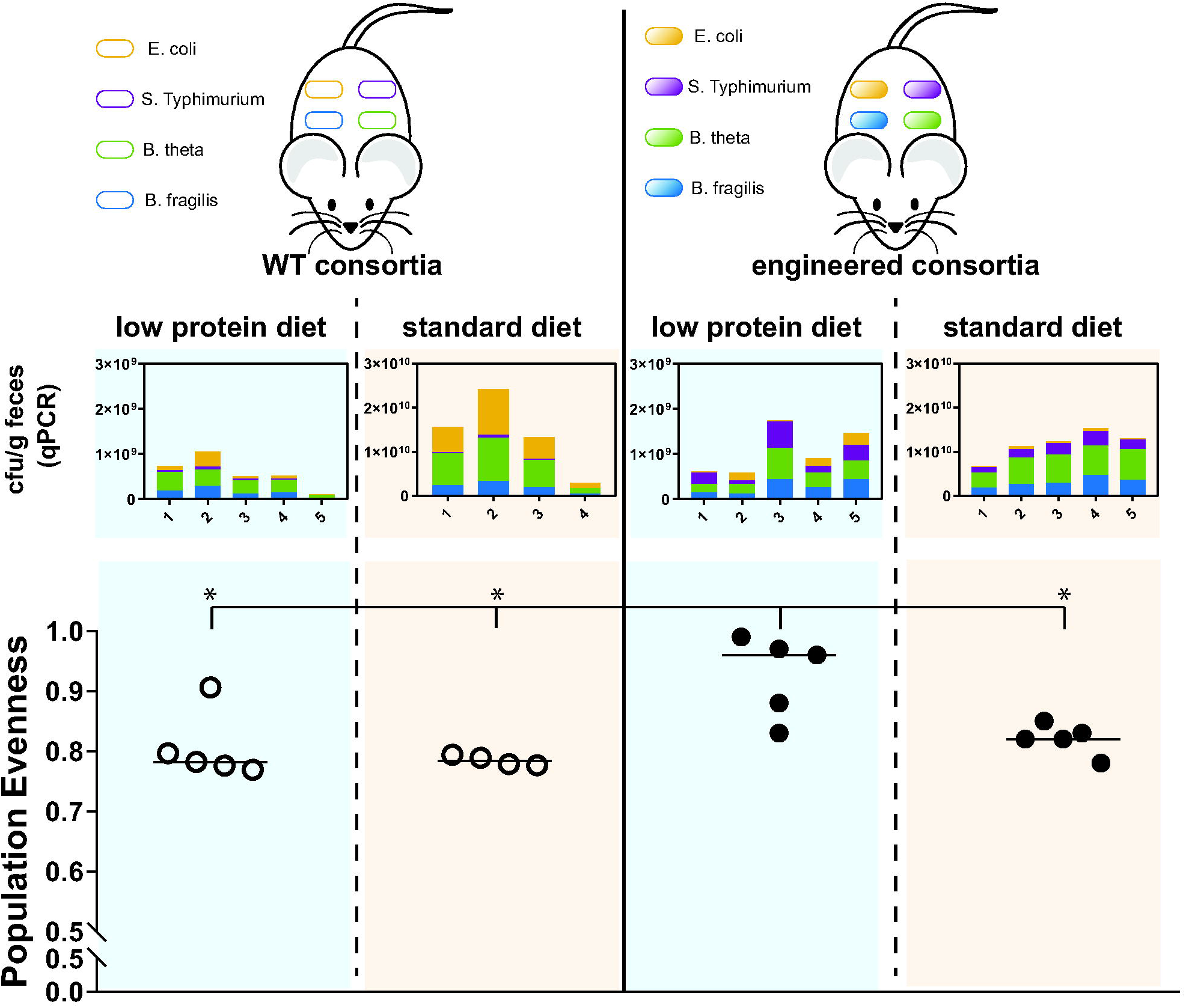
Consortia engineering increases population evenness in the mammalian gut in a diet dependent manner. Four groups of germfree mice (n=5, except second group from the left which is n=4) were either fed low protein diet or standard diet, and inoculated with either the engineered or WT bacterial consortia. Fecal samples 10 days post-inoculation were analyzed via strain-specific qPCR to assess concentrations of each consortia species. Population evenness of consortia was calculated. Bar indicates Median. Mann-Whitney test showed significantly increased population of the engineered consortia in mice that were fed low protein diet compared to the consortia in the three other groups (p-values: 0.024, 0.032, 0.015). yellow: *E. coli*; purple: S. Typhimurium; green: *B. thetaiotaomicron;* blue: *B. fragilis.*

We observed consistently higher abundances of all strains in both engineered and WT consortia in mice that were fed with the standard diet. For the engineered consortium, species abundances were higher by a factor of approximately 3 for *E. coli,* 8 for S. *Typhimurium,* 16 for *B. thetaiotaomicron,* and 11 for *B. fragilis* in mice fed a standard versus a low protein diet. In the case of the WT consortium, S. *Typhimurium, B. thetaiotaomicron* and *B. fragilis* concentrations were similarly higher on standard chow (fold changes of approximately: 10, 22, 13 respectively), and WT *E. coli* concentrations were considerably higher (~50-fold higher).

Although overall strain abundance was higher in mice that were fed with standard chow, the engineered consortium in mice that were fed with a low protein diet showed significantly greater population evenness (Mann-Whitney test; p-values: 0.02, 0.03, 0. 02). These findings are consistent with our *in vitro* results showing the greatest population evenness in the engineered consortium in conditions with low concentrations of the cross-fed amino acids. We also observed a trend toward greater population evenness in the engineered consortium in mice fed standard chow compared to the WT consortium in mice regardless of diet, although this trend was not statistically significant. Interestingly, in mice that were fed with a low protein diet, S. Typhimurium grew about 8-fold better in the engineered consortium compared to the WT consortium. This finding is consistent with our *in vitro* results, which indicated that engineered S. Typhimurium benefits most from the consortium cross-feeding effects. We observed the same trend in mice that were fed with standard diet, albeit to a lesser extent

## Discussion

We engineered inter-species amino acid cross-feeding in a bacterial consortium and demonstrated that this increased population evenness in environments of varying complexity. The ability of an ecosystem to maintain evenness, an important component of species diversity, in the face of different environments or perturbations to the same environment is a critical component to its robustness or limiting dominance of some ecosystem members at the expense of others. We showed that pre-existing antagonistic interactions in a bacterial consortium can be overcome by engineered beneficial interactions (Figure 2). The engineered consortium exhibited increased population evenness *in vitro,* particularly in nutrient sparse conditions when crossfeeding was favored (Figure 3).

We found that the engineered consortium also showed increased population evenness in the gnotobiotic mouse gut (Figure 4). Gnotobiotic mice provide a tractable model for the mammalian gut environment, maintaining the spatial organization and metabolic composition of the host. However, a limitation is that gnotobiotic mice lack the myriad of bacterial species present in a naturally occurring commensal microbiota (38). Despite this limitation, our results provide evidence that engineered bacterial cross-feeding behavior can function in a complex environment and maintain increased population evenness. Notably, while the engineered consortium attained the highest population evenness *in vitro* in conditions lacking the cross-fed amino acids, this was at the expense of overall biomass. However, in the gnotobiotic mouse gut, the engineered consortium attained about the same biomass as its WT counterpart in mice receiving the same diet. Given that the strains in the consortium all originated from the mammalian gut, these findings reinforce the fact that *in vitro* conditions may not mimic important aspects of the host environment such as availability of complex nutrients and chemical environments in the gut as well as differing spatial distribution of bacterial species, i.e., in large versus small bowel or luminal versus mucosal surfaces. An interesting direction for future work would be to better characterize the niches of the consortium member in the gut, such as through *in situ* localization assays, and determine whether engineering alters spatial localization and other niche behavior.

Engineering diverse, multi-species consortia presents interesting challenges while holding promise for substantial improvements for applications in human or animal health and bioproduction. Bacteria from naturally occurring ecosystems will have pre-existing interactions, which are often competitive (4–6). In a given bacterial consortium that is dominated by antagonistic interactions, synthetically introduced positive interactions need to overcome these antagonistic interactions in order to cause a net beneficial effect. We found that in our consortium there was considerable bias toward naturally occurring competitive interactions. Thus, any engineering of these organisms had to overcome this inherent negative bias. We engineered bacteria to overproduce metabolites at different concentrations from low *(S.* Typhimurium, His) to high *(B. fragilis,* Arg) (Figure 1). This enabled us to assess the effects of various levels of overproduction in a single consortium. *B. fragilis* overproduction achieved the most obvious beneficial interactions and its knockdown *in vitro* had the most profound effect on consortium growth (Figure 2). Overproduction by the other strains led to reduction of pre-existing antagonistic interactions, but not to the extent of producing fully beneficial interactions.

Many studies on engineered cross-feeding in microbial consortia have focused on knockouts and complementary metabolism but not overproduction (18, 21). Our findings suggest that relatively high overproduction rates are required in a consortium where multiple members cross-feed from a single or small number of overproducers. We did not observe reduced growth of the overproducing strains, with the exception of *B. thetaiotaomicron,* which showed a 3-fold reduction in growth (Figure S5). Our findings have implications for naturally occurring systems, and merit further study to understand the extent to which robust cross-feeding has evolved to depend on multiple producers versus high output single producers.

We observe that our engineered consortium exhibits increased evenness at the cost of reduced fitness; the engineered consortium grows at ~10-fold lower concentration than the WT counterpart (Figure 2). This is likely in part due to the fact that some of the bacteria overproduce less than is required by the other members (Figure 1). This imbalance may also make the consortium susceptible toward cheaters; in fact *E. coli* and S. Typhimurium could be viewed as “moochers” that benefit most from *Bacteroides spp.* production but do not contribute much themselves. Future work could seek to optimize the balance of overproduction and consumption rates of the consortium members to improve overall consortium fitness while preserving population evenness, using approaches such as rationally engineering overproduction pathways and amino acid uptake transporters in combination with directed evolution via turbidostat.

Our finding that metabolite cross-feeding can lead to increased population evenness, which is an important aspect of species diversity, may have implications for understanding naturally occurring microbial ecosystems as well as engineered communities for biotechnology applications. Indeed, prior work has shown that in a methanotrophic bacterial consortium, amino acid cross-feeding helps to maintain species diversity (15). Species diversity is clearly important in the robustness of microbial ecosystems; an ecosystem with greater genetic diversity as a whole has the potential to better survive in changing environments (e.g. levels of cross-feed nutrients for our consortium). The gut microbiota constitutes a complex host-microbial ecosystem and maintaining species diversity in this context has important implications for human health. Our internal microbial ecosystems undergo a variety of alterations due to factors including normal physiologic changes such as exercise or pregnancy, pathologic states due to disease, modifications to our diets, and medical interventions. Thus, mechanisms such as cross-feeding that promote maintenance of species diversity in the human microbiome in the face of perturbing factors may be critical to maintaining host function. From the perspective of an ecosystem, there is a long-term evolutionary fitness advantage to maintaining species diversity, which may be supported by naturally occurring metabolite cross-feeding mechanisms such as those we introduce artificially here.

## Acknowledgement

We thank Laurie Comstock for providing *B. fragilis* 638R and helpful advice on *Bacteroides* handling. This project was funded by the Defense Advanced Research Program Agency (DARPA BRICS HR0011-15-C-0094), NIH T32 HL007627 (to T.G.), the Harvard Digestive Diseases Center P30 DK034854 and The Wyss Institute for Biologically Inspired Engineering. D.T.R. is supported by a Human Frontier Science Program Long-Term Fellowship and a National Health and Medical Research Council (NHMRC) RG Menzies Early Career Fellowship from the Menzies Foundation Australia.

## Methods

### Auxotroph engineering

For auxotroph generation in the *E. coli* NGF-1 strain we introduced multiple knockouts using sequential P1 transduction (39) from the Keio knockout collection (40). Flip-out of kanamycin cassettes was done using pCP20 (41). In brief, for P1 transduction we prepared phage by diluting an overnight culture of the donor strain 1:100 LB with 0.2% glucose, 5 mM CaCl_2_ and 25 mM MgCl_2_ and incubated for 1-2 hours at 37 °C until slightly turbid. We then added 40 μL P1 lysate and continued growth for 1-3 h at 37 °C while shaking until lysed. Lysate was then filtered with a 20 μm sterile filter and stored in the fridge. For transduction, we harvested 2 mL overnight culture of recipient strain and re-suspended in 2 mL LB with 5 mM CaCl_2_ and 100 mM MgSO_4_. We then mixed 100 μL donor lysate with 100 μL recipient, incubated 30 min at 37 °C and added 200 μL sodium citrate (1 M, pH 5.5) and 1 mL LB and incubated for another 1 hr at 37 °C. Cells were harvested, re-suspended in 100 μL LB with 100 mM sodium citrate and plated on LB Kan plates (75 μg/mL). The transduced kanamycin cassette was then removed using pCP20 according to protocol. We transformed pCP20 via electroporation and transformants were selected on LB agar plates supplemented with 100 μg/mL carbenicillin grown at 30 °C. Single colonies were re-streaked on LB without drugs and incubated for 10 hours at 42 °C. From there, single colonies were re-streaked on LB plates without drugs and grown overnight at 37 °C. Colonies were checked for Carbenicillin, and Kanamycin sensitivity and further confirmed via PCR at respective loci. This procedure was repeated until all knockouts were introduced.

Engineering of S. Typhimurium LT2 required generation of single knockout strains that contained pKD46 integrated into the genome, which allowed for linear DNA integration using lambda red recombination (41). We then introduced the knockouts into the S. Typhimurium strain through sequential P22 transduction and pCP20 flipout analogous to *E. coli* engineering. Single knockout strains were generated by PCR amplifying a Kanamycin resistance cassette from pKD13 generating linear fragments that contained upstream and downstream homology to the gene of interest and the kanamycin cassette with FRT sequences. Fragments were introduced via electroporation and selected on LB agar plates supplemented with 50 μg/mL Kanamycin. Sequential P22 transduction and pCP20 flip-out was essentially performed as described above for P1 transduction but lysis was done overnight.

For knockout generation of both *B. thetaiotaomicron* and *B. fragilis,* we used pExchange KO vectors as described (42). Briefly, we introduced 750 bp flanking regions for genes of interest adjacent to each other into the vector. The vector contains an erythromycin resistance positive marker and a thymidine kinase as counter selection marker. Cloning was done in pir+ *E. coli* strains and vectors were transferred to MFDpir for conjugation (43). Conjugation was done according to protocol with minor changes. In brief, five drops of overnight culture of *E. coli* donor was inoculated in LB supplemented with 300 μM Diamino pimelic acid (DAP) and five drops of recipient overnight culture was inoculated in 50 mL basal media. Both cultures were grown for about 2 hr (E. *coli* aerobically, *Bacteroides spp.* anaerobically) until *E. coli* culture was well turbid and *Bacteroides* culture just slightly turbid. Subsequently, 9 mL recipient and 3 mL donor were combined and spun down for 10 min at 4000 rpm together. The pellet was re-suspended in 100 μL fresh basal media with 300 μM DAP and pipetted on basal media agar plates without cysteine and supplemented with 300 μM DAP. The cells were incubated at 37 °C aerobically face up for up to 20 hr, scraped off and re-suspended in 10% glycerol. Dilutions were plated on basal-agar plates supplemented with 10 μg/mL erythromycin and incubated at 37 °C anaerobically for 2-3 days. Single colonies were re-streaked in the presence of erythromycin and grown for another 2 days. 10 single colonies were inoculated in basal media without drug and grown overnight. 500 μL of each culture was mixed, spun down and re-suspended in 10% glycerol. We then plated different dilutions on basal media plates supplemented with 5-fluoro-2-deoxy-uridine (FuDR) (200 μg/mL) and incubated at 37 °C anaerobically for 3 days. Knockouts were verified via PCR. This procedure was repeated multiple times to obtain the multiple auxotroph strains.

### Overproducer selection

Overproducers were generated by selecting for mutants that could grow on minimal media agar plates supplemented with anti-metabolites (E. *coli:* 5 mg/mL Norleucine for Met overproduction; S. Typhimurium: > 0.7 mg/mL beta-(2-thiazolyl)-DL-alanine for His overproduction; *B. thetaiotaomicron:* 50 μg/mL 4-methyl tryptophan for Trp overproduction; *B. fragilis:* 80 μg/mL Canavinine for Arg overproduction). Single colonies that showed halos were re-streaked and overproduction was measured using a bioassay. In brief, for screening of overproducing mutants the isolated strains were grown overnight at 37 °C shaking aerobically (for *E. coli* and S. Typhimurium) or anaerobically without agitation (for *Bacteroides spp.).* Supernatant was harvested, diluted 1:1 with fresh media, and *E. coli* auxotrophs were inoculated and their growth was recorded after 24 hr. For *E. coli* NGF-1 overproducers, we used a S. Typhimurium auxotroph instead, since its colicin production prevented the *E. coli* biosensor from growing. Confirmed overproducers were further quantified using LC-MS.

### LC-MS for Overproduction Measurements

To quantitate amino acid levels in overproducer supernatants, a standard curve was obained using freshly prepared amino acid standards dissolved in growth media (1mM, 500 uM, 100 uM, 50 uM, 10 uM of L. Methionine, L/Histidine, L-Tryptophan, L-Arginine each). To prepare for HPLC-MS analysis, 0.5 mL sample or standard were added to 1.5 mL ice-cold methanol and incubated on ice for 10 min. The mixture was centrifuged for 5 min at 15,000 rpm and 500 μL supernatant was vacuum concentrated and re-suspended in 50 μL methanol. Samples were kept on ice or at 4°C. HPLC-MS analysis of standards and extracts was carried out using an Agilent 1260 Infinity HPLC system equipped with an Agilent Eclipse Plus C18 (100 × 4.6 mm, particle size 3.5 mm, flow rate: 0.3 mL/min, solvent A: dd.H_2_O/0.1% (v/v) formic acid, solvent B: acetonitrile, injection volume: 4 mL) connected to an Agilent 6530 Accurate-Mass Q-TOF instrument. The following gradient was used (time/min, %B): 0, 0; 0.5, 0; 14, 100; 19, 100; 20, 0, 25, 0. The mass spectrometer was operated in positive mode and the autosampler was kept at 4°C. After HPLC-MS analysis, extracted ion current (EIC) peaks were automatically integrated using the MassHunter Workstation Software (version: B.07.00). A plot of peak area versus amino acid concentration was used to generate a linear fit.

### Sequencing

Bacterial cultures were prepared in rich media (basal for *Bacteroides spp.* and LB for *E. coli* and S. Typhimurium). Genomic DNA (gDNA) extraction was performed using the Wizard Genomic DNA Purification Kit (Promega) according to protocol. The extracted gDNA was sheared using Covaris DNA Shearing, and the library was prepared using Kapa Biosystem DNA Hyper Prep NGS Library (Dana Faber Core MBCFL Genomics). Sequencing was performed on the Illumina MiSeq instrument, with the 150 bp paired End (PE150) reagents. Sequences were analyzed for SNPs using Geneious software and published genome sequences *(E. coli:* CP016007.1; S. Typhimurium: NC_003197; *B. thetaiotaomicron:* AE015928; *B. fragilis:* NC_016776).

### Growth and Media Conditions

All basal media (20 g/L proteose peptone, 5 g/L yeast extract, 5 g/L sodium chloride, 1 g/L L-cysteine, 0.5% glucose, 5 mg/mL potassium phosphate, 5 mg/L hemin) and co-culture media (M9 salts (0.2 g/L Na_2_HPO_4_, 90 mg/L KH_2_PO_4_, 30 mg/L NH_4_Cl, 15 mg/L NaCl), 1 mM MgSO_4_, 10 μg/mL heme, 0.1 mM CaCl_2_, 1 μg/mL Niacinamide, vitamin B12 and thiamine, 400 μg/mL L-cysteine, 0.3% bicarbonate buffer, 2.5 ng/mL vitamin K, 2 μg/mL FeSO_4_*7H_2_O and carbon sources and amino acid supplementation as described in Results) were pre-incubated for at least 24 hr anaerobically before use. In initial experiments we used 0.5% starch, which we later optimized to 0.2% (resulting in lower viscosity); the observed growth differences were not statistically significant between those two conditions (Figure S7). *Bacteroides spp.* were inoculated from glycerol stock into basal media, grown overnight and 400 μL was inoculated in 5 mL basal and grown 2 hr anaerobically. Cells were spun down, washed twice in PBS and diluted in growth media as described for each experiment in Results. *E. coli* and S. Typhimurium were inoculated from glycerol stock into LB and grown overnight at 37 °C while shaking. 100 μL of culture was then inoculated into pre-incubated LB and grown anaerobically for 2 hr, diluted, washed in PBS and diluted into co-culture media as described.

### Multiplex qPCR

We designed strain specific primer/probe-fluorophore pairs according to IDT protocol (Table S1). We chose strain specific genes by multiple genome alignment between the strain of interest, the other consortia members and closely related strains using Mauve (44). Multiplex qPCR was used to quantify each strain in co-culture by using a standard curve obtained by plating late log phase cultures grown in rich media. In brief, each strain was grown from overnight culture for ~5 hours until about OD of 1. Cells were then counted by plating. Cultures were mixed, diluted and frozen at −80 °C for use as standard curve. Samples were diluted 1:10 in ddH_2_O and snap-frozen in liquid nitrogen and stored at least overnight at −80 °C. Growth curve and sample were both thawed together and prepared in a 5 μL Primetime Mastermix (IDT) with 1 μL Primer/Probe mixture (final concentrations: 100 nM for primers and 50 nM for probes). The qPCR was run with the following program: 20 min at 98 μC (to boil the cells and denature gDNA), followed by 40 cycles of 60 °C and 98 °C.

### Statistical inference of interactions within consortia

We adapted our previously published statistical method for inferring a dynamical systems model from microbial time-series data (33, 34). Briefly, our model assumes the rate of change over time of each microbe in the ecosystem is related to its own abundance as well as abundances of the other microbes in the ecosystem. Our model is fully Bayesian and infers the posterior probability distribution over the qualitative network of microbe-microbe interactions (e.g., probability of an edge being present in the network) as well as the quantitative strengths of the interactions present. As we have previously described (33), the qualitative network can be interpreted in terms of Bayes factors (BF), a standard Bayesian alternative for hypothesis testing that quantifies the evidence for one model over another. For our method, the BF on network edges quantifies the evidence for existence of the interaction versus absence of the interaction. A standard interpretation of BFs is: BF between 3 and 10 = substantial evidence, BF > 10 = strong evidence.

To be precise, our model is based on discrete time stochastic generalized Lotka Volterra (gLV) dynamics:

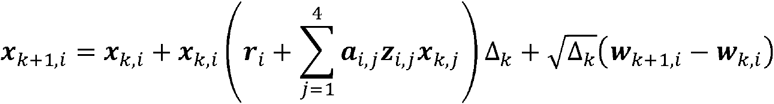

where ***x**_k_,i* is the abundance of species *i* at time point *k; **r**_t_* is the growth rate of species *i; **a**_i,j_* is the interaction coefficient and ***z**_i,j_* is a binary variable indicating the presence or absence of an edge (interaction) between species *j* and *i*; Δ_*k*_ is the time difference between timepoints *k* + 1 and *k*·, and ***w**_k,i_* is a Brownian motion term. The Brownian motion has variance 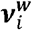. We place prior probability distributions on the variables defined above:

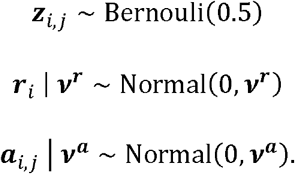

Note that our prior probability distribution for the indicator variable, ***z**_i,j,_* for presence or absence of interactions, expresses maximum uncertainty, e.g., no *a priori* assumption about the presence or absence of an interaction. We place the following prior probability distributions on the variance parameters in our model:

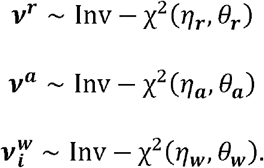

In our model *η_**r**_η_**a**_η_**w**_* = .1 with *θ_**r**_* = .01, *θ_**a**_* = 10^4^, and *θ_**w**_* = 10^4^, specifying relatively diffuse (uninformative) priors. We perform inference using a simplified version of our previously described Markov Chain Monte Carlo (MCMC) algorithm that omits clustering (34), with 5,000 iterations (1,000 burn in iterations.)

The gLV model described above allows for only pairwise interactions, so we also tested an extended model with higher order interactions. Specifically, we included 3^rd^ order or 4^th^ order interactions, i.e.,

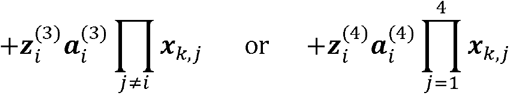

appearing on the right-hand side of the gLV dynamics equation above. The superscripts (3) and (4) denote higher order indicator variables 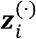 and interaction coefficients 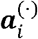 for orders 3 and 4 respectively. However, when we applied the model to our data, pairwise interactions dominated, and the indicator variables associated with the higher order interactions had Bayes factors near zero. From these analyses, we can conclude that the presence of higher order interactions was not supported by our data. However, it is possible that such interactions actually exist in the ecosystem but that our data is insufficient for rigorous inference using our statistical method.

### Calculation of Population Evenness (Pielou’s Evenness)

Species Diversity (Pielou’s evenness) was calculated according to the given formula:

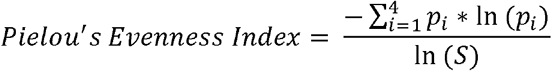

Where *p_i_*, refers to the population ratio of a given strain in the consortium of four strains. *S* is the number of species.

### *In vivo* Experiments

Adult (6-8 weeks) male Swiss Webster germ free mice bred in house at the Massachusetts Host-Microbiome Center were used. Animals were fed either on standard chow in the facility for the entire experiment, or on low-protein diet (3% custom diet, envigo, doubly irradiated) beginning 10 days prior to the experiment and continuing for its duration. To prepare bacteria for gavage, we grew each strain to mid-log phase, plated for counting and snap-froze aliquots. For gavage, aliquots were thawed, spun down, combined to achieve concentrations of approximately 10^7^ per bacteria per gavage, and re-suspended in 200 μL 1x PBS with 0.05% L-cysteine for gavage. After gavage, mice were transferred to Optimice cages and maintained gnotobiotic for 10 days. Fecal samples were collected prior gavage and at 10 days, and snap-frozen for storage at −80 °C.

### Molecular Analysis of *in vivo* Samples

DNA was extracted from fecal samples using the Zymobiomic 96 DNA Kit with the following modification: we omitted the silicon-ATM-HRC wash. Cells were lysed in a bead beater at speed 20 for 10 min, plates were turned and lysed for another 10 min at the same speed. We added an additional 3 min incubation step for Binding Buffer and additional 5 min incubation steps when transferring to Zymo-Spin-I-96-Z plates. Elution was done in 50 μL ZymoBIOMIC DNase/RNase Free Water.

Direct multiplex probe-based qPCR was done on extracted DNA samples as described above. For standard curves, we used plated overnight cultures spiked into germfree fecal samples and extracted them as described above.

## Supplemental Material

**Figure S1** Growth response. Each auxotroph was grown in media supplemented with varying concentrations of one amino acid and saturating concentration of the two others. Depicted is the average of three biological replicates; error bars indicate standard deviation. A sigmoidal curve was fit using GraphPad Prism 8.

**Figure S2** Log fold ratios (LogR) trajectories in WT consortium in all four conditions compared to condition one in Figure 2. Legend: Ec = *E. coli,* ST = S. Typhimurium, BT = *B. theta,* BF = *B. fragilis.*

**Figure S3** Log fold ratios (logR) trajectories in engineered consortium in all four conditions compared to condition one in Figure 2. Legend: Ec = *E. coli,* ST = S. Typhimurium, BT = *B. theta,* BF = *B. fragilis.*

**Figure S4** Growth rates, Bayes Factors (for microbial interactions), and Microbial Interaction Strengths for unengineered (A) and engineered (B) consortia learned from *in vitro* growth data. The inferred growth rates for the engineered consortia are smaller by about an average factor of 3 for any species. The Bayes Factors quantify the evidence for an interaction being present verses being absent (Bayes Factor between 3 and 10 = substantial evidence, Bayes Factor > 10 = strong evidence). The inference is highly confident in 7 of the 12 unenginereerd consortia interactions. Note that an infinite Bayes Factor means that the edge was included in every single step when learning the posterior distribution. For the engineered consortia two of the 12 interactions exist with at least substantial evidence with 5 of the interactions having very low evidence for the edge being present in the model but still greater than evidence for no interaction (Bayes Factor of 1 is the scenario where evidence for and against an interaction being present are equal). The interaction strengths for the engineered consortia are of higher magnitude when compared to the unengineered consortia. This occurs because the carrying capacity of the unengineeered consortia is about 1.5 orders of magnitude larger than the engineered consortia and microbe-microbe interactions are inversely proportional to the carrying capacity in a gLV model.

**Figure S5** Growth of WT and engineered strains after 24 hr, at which point supernatant was collected for cross-feeding experiment (see Figure 1). Overproduction does not affect growth with the exception of *B. theta* (3-fold reduction). Shown are three biological replicates with median indicated as horizontal line.

**Figure S6 Metabolite overproduction and WT and engineered strains**. Amino acid production was measured using LC-MS. The average of three biological replicates is shown.

**Figure S7 Growth after 24 hr in media supplemented with 0.5% or 0.2% starch**. Growth did not differ significantly for any of the strains (Mann-Whitney test, p-values > 0.1).

**Table S1 qPCR probes and primers**.

