## Supplemental Material for "Engineered inter-species amino acid cross-feeding increases population evenness in a synthetic bacterial consortium"

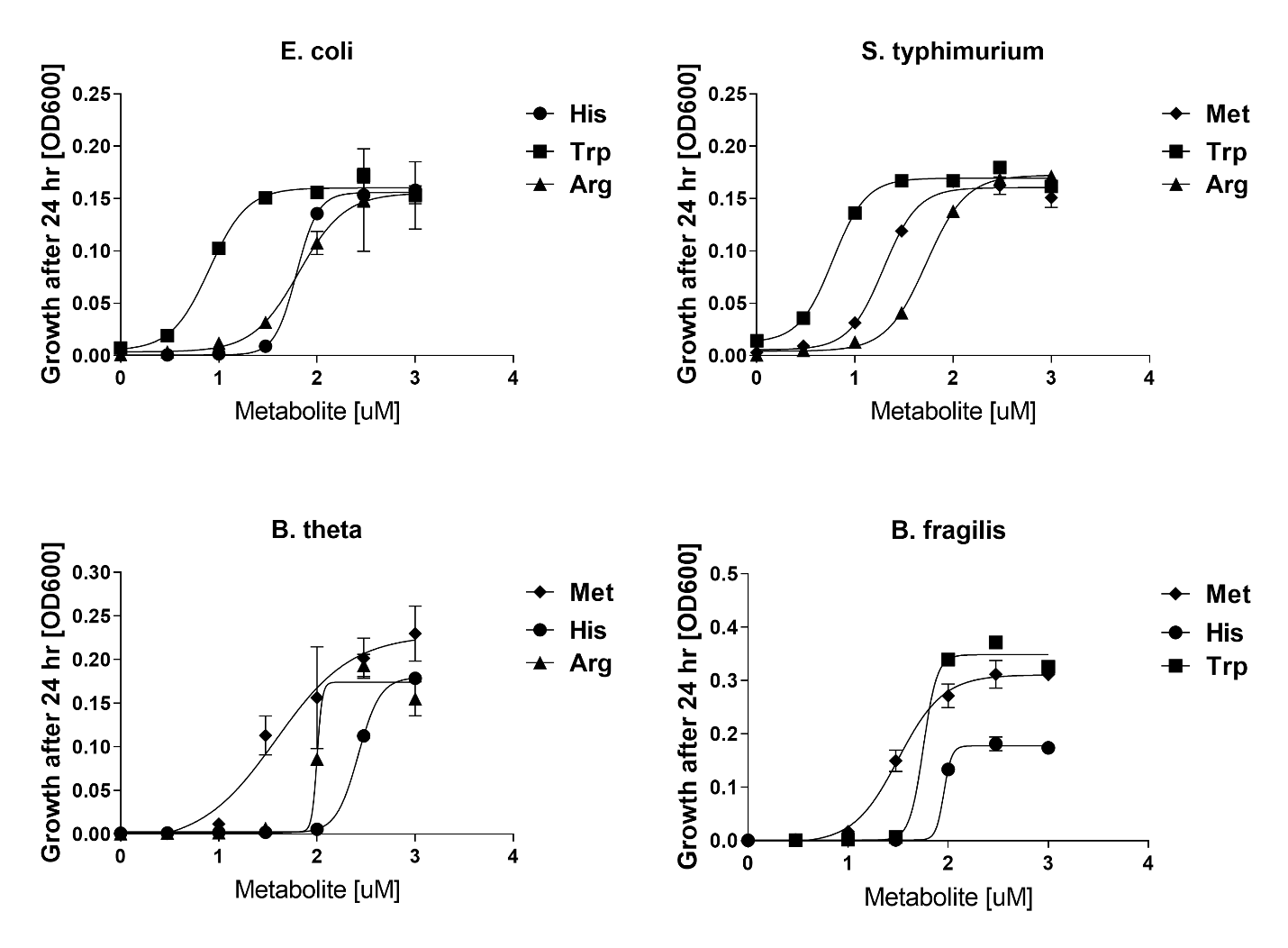

**Figure S1** Growth response. Each auxotroph was grown in media supplemented with varying concentrations of one amino acid and saturating concentration of the two others. Depicted is the average of three biological replicates; error bars indicate standard deviation. A sigmoidal curve was fit using GraphPad Prism 8.

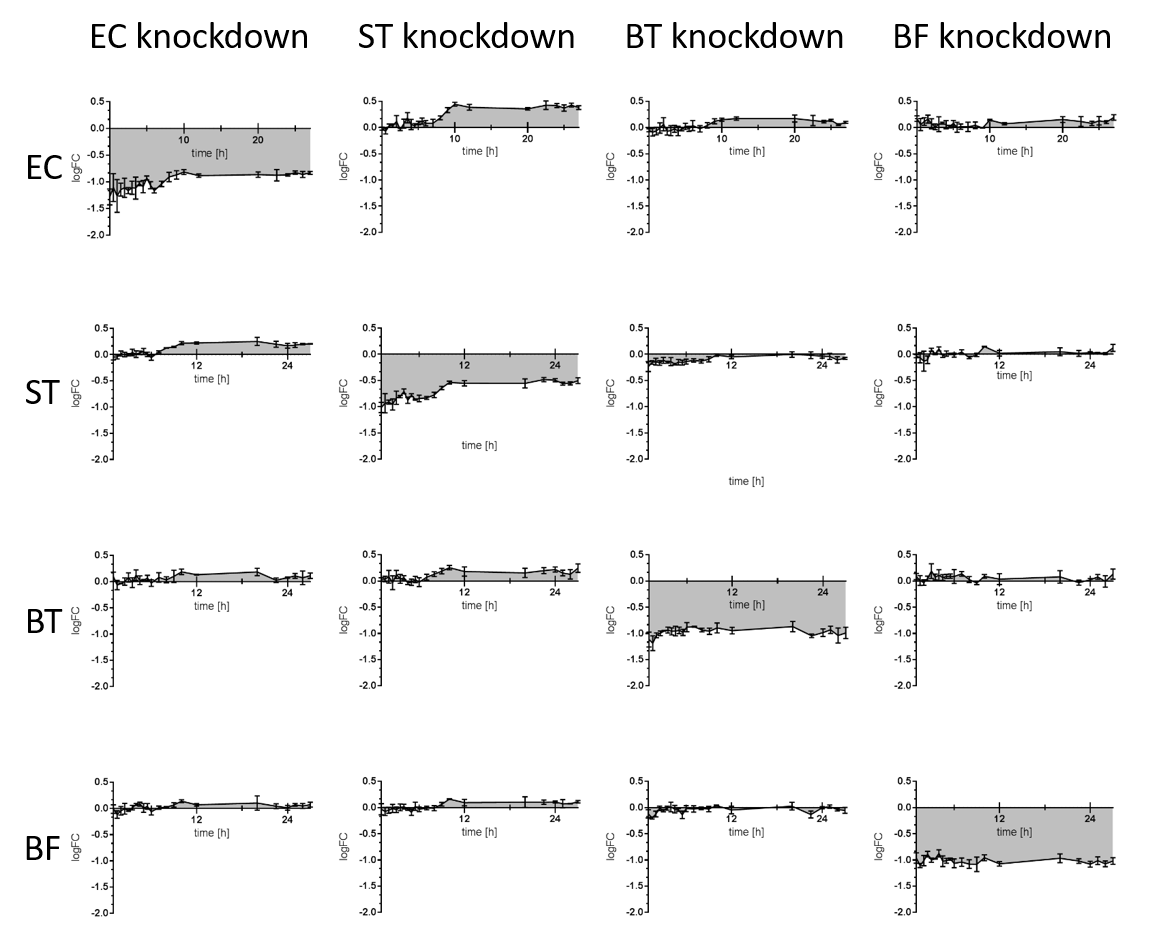

**Figure S2** Log fold ratios (LogR) trajectories in WT consortium in all four conditions compared to condition one in Figure 2. Legend: Ec = *E. coli*, ST = *S.* Typhimurium, BT = *B. theta*, BF = *B. fragilis*.

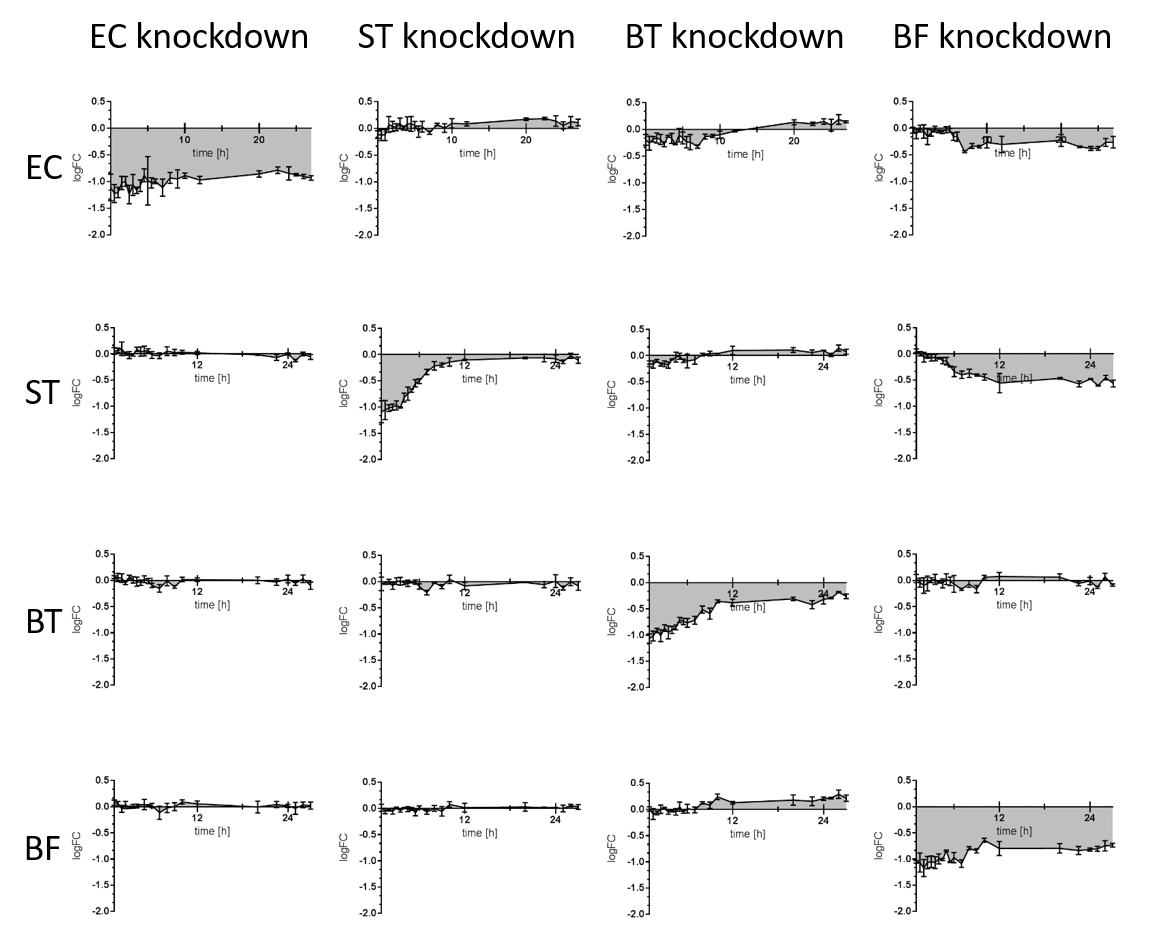

**Figure S3** Log fold ratios (logR) trajectories in engineered consortium in all four conditions compared to condition one in Figure 2. Legend: Ec = *E. coli*, ST = *S.* Typhimurium, BT = *B. theta*, BF = *B. fragilis*.

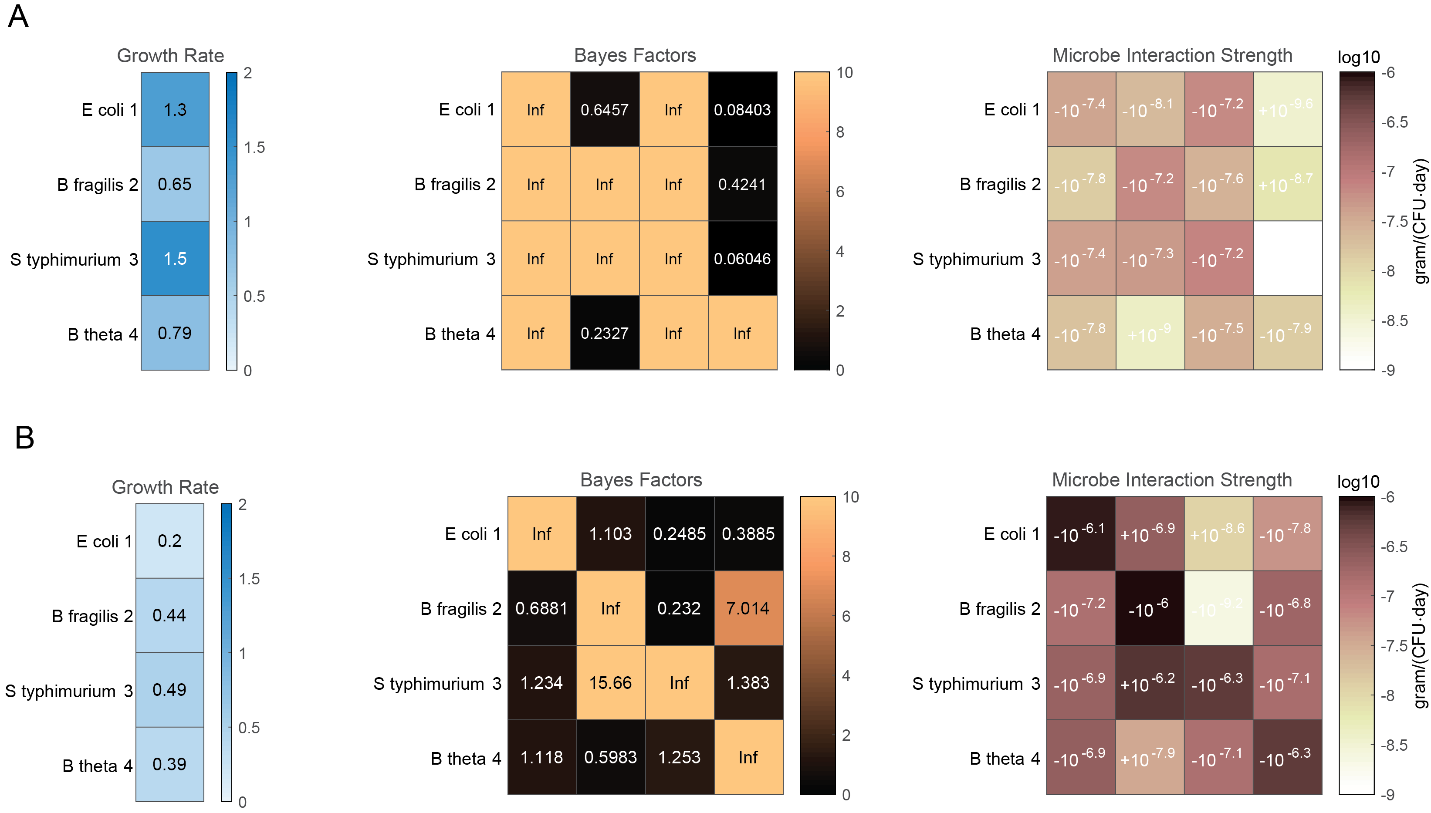

**Figure S4** Growth rates, Bayes Factors (for microbial interactions), and Microbial Interaction Strengths for unengineered (A) and engineered (B) consortia learned from *in vitro* growth data. The inferred growth rates for the engineered consortia are smaller by about an average factor of 3 for any species. The Bayes Factors quantify the evidence for an interaction being present verses being absent (Bayes Factor between 3 and 10 = substantial evidence, Bayes Factor > 10 = strong evidence). The inference is highly confident in 7 of the 12 unenginereerd consortia interactions. Note that an infinite Bayes Factor means that the edge was included in every single step when learning the posterior distribution. For the engineered consortia two of the 12 interactions exist with at least substantial evidence with 5 of the interactions having very low evidence for the edge being present in the model but still greater than evidence for no interaction (Bayes Factor of 1 is the scenario where evidence for and against an interaction being present are equal). The interaction strengths for the engineered consortia are of higher magnitude when compared to the unengineered consortia. This occurs because the carrying capacity of the unengineeered consortia is about 1.5 orders of magnitude larger than the engineered consortia and microbe-microbe interactions are inversely proportional to the carrying capacity in a gLV model.

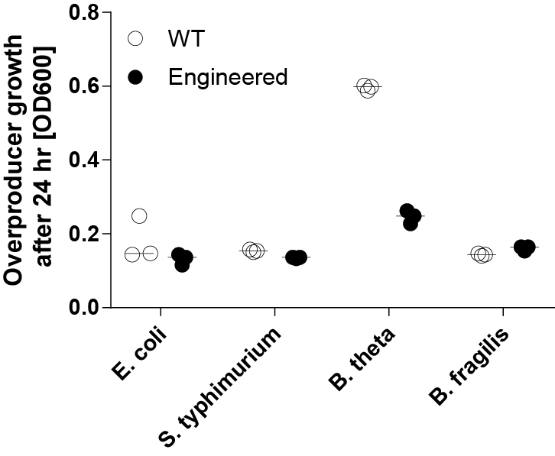

**Figure S5** Growth of WT and engineered strains after 24 hr, at which point supernatant was collected for cross-feeding experiment (see Figure 1). Overproduction does not affect growth with the exception of *B. theta* (3-fold reduction). Shown are three biological replicates with median indicated as horizontal line.

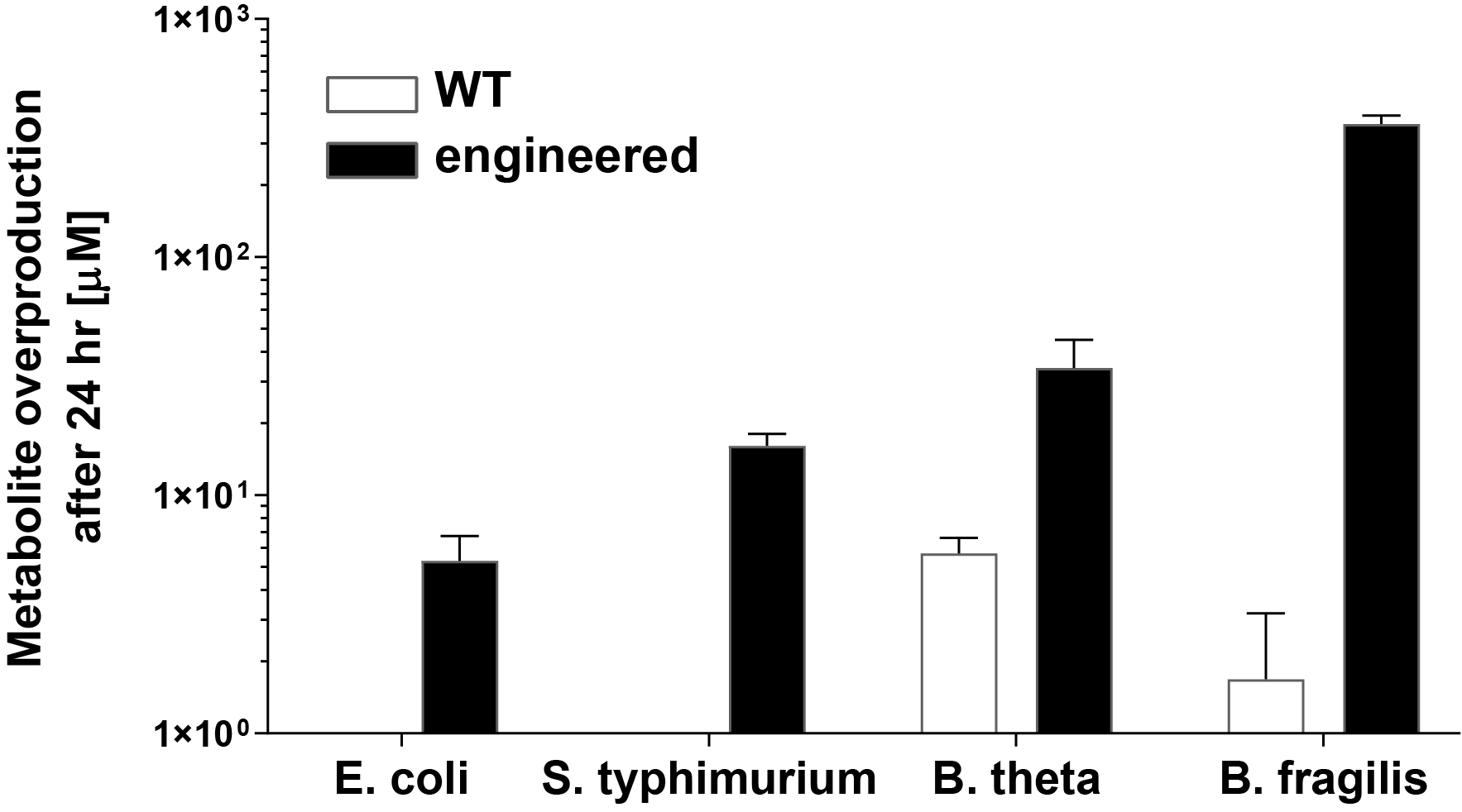

**Figure S6 Metabolite overproduction and WT and engineered strains.** Amino acid production was measured using LC-MS. The average of three biological replicates is shown.

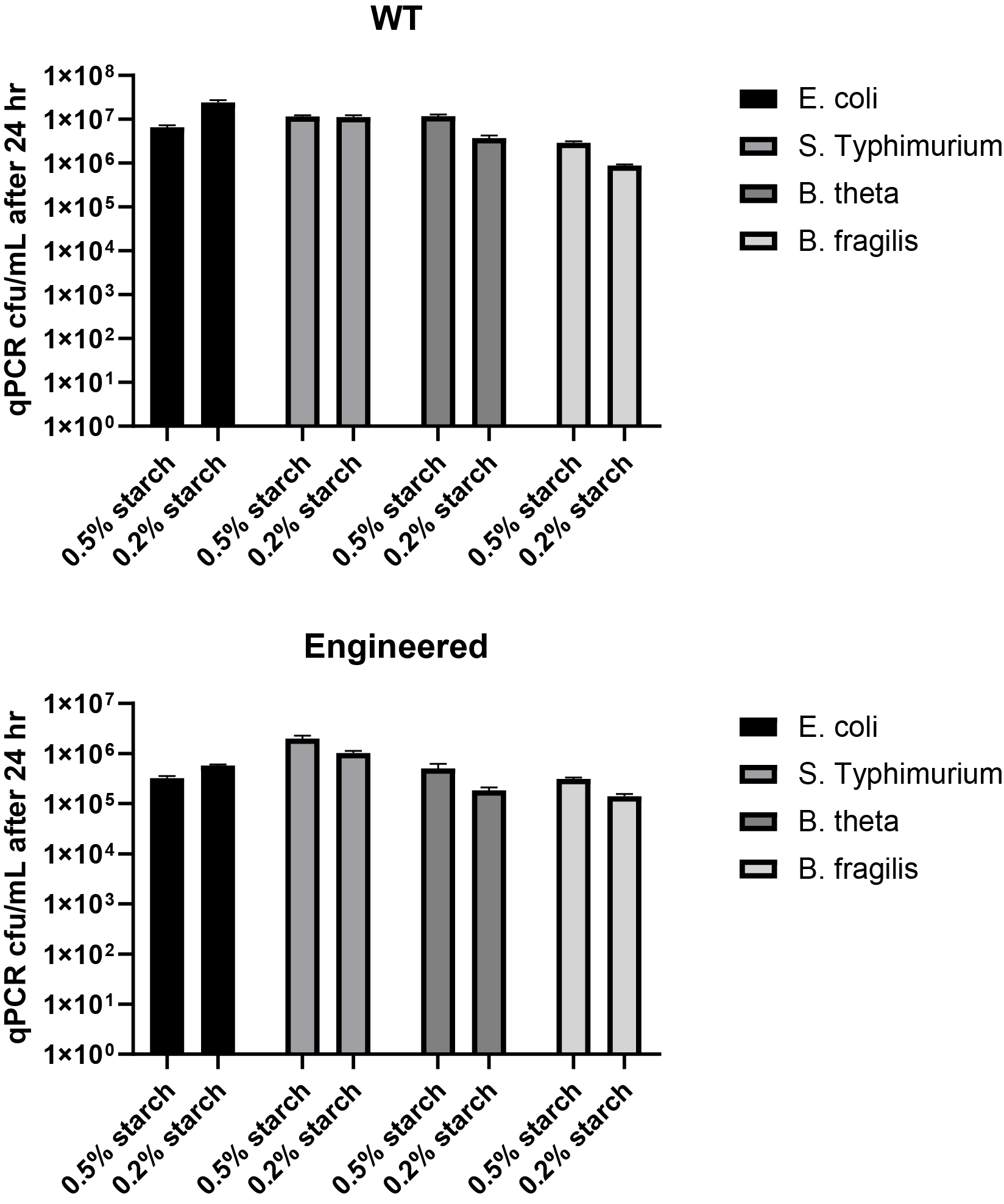

**Figure S7 Growth after 24 hr in media supplemented with 0.5% or 0.2% starch.** Growth did not differ significantly for any of the strains (Mann-Whitney test, p-values > 0.1).

**Table S1 qPCR probes and primers.**

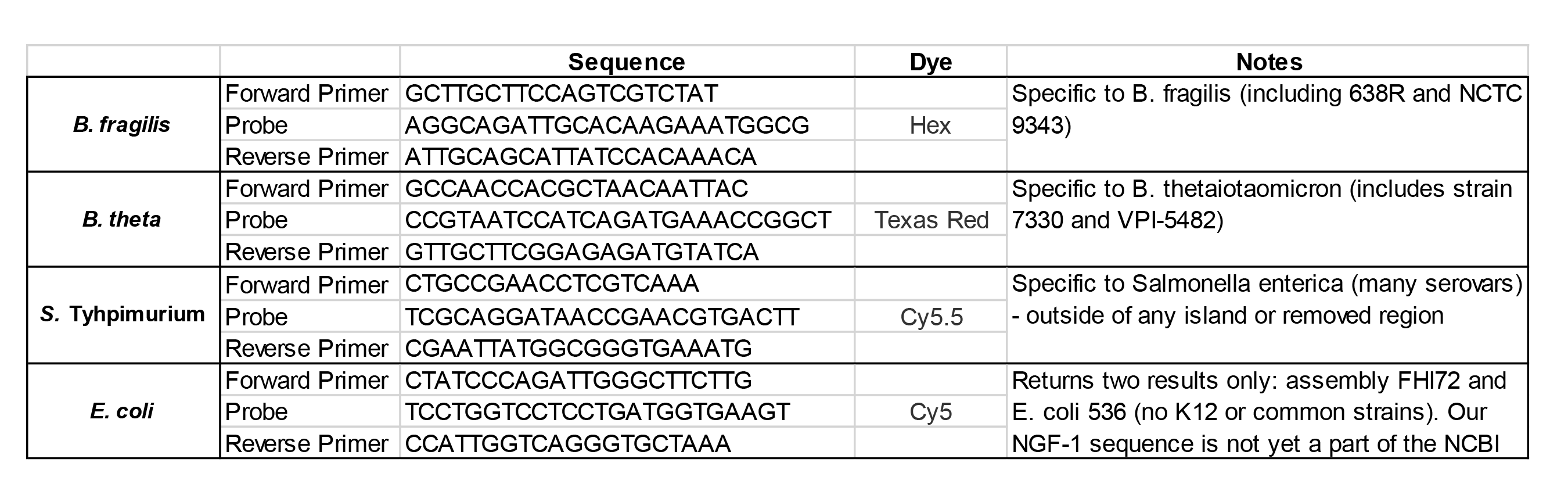
